# Loss of SPNS1, a lysosomal transporter, in the nervous system causes dysmyelination and white matter dysplasia

**DOI:** 10.1101/2024.05.29.596535

**Authors:** Yoshinobu Ichimura, Yuki Sugiura, Yoshinori Katsuragi, Yu-Shin Sou, Takefumi Uemura, Naoki Tamura, Satoko Komatsu-Hirota, Takashi Ueno, Masato Koike, Satoshi Waguri, Masaaki Komatsu

## Abstract

Protein spinster homolog 1 (SPNS1) is a lysosomal transporter of lysophospholipids and sphingosine, which has recently been identified to be mutated in patients with neurodegeneration. However, its physiological role, especially in the nervous system, remains largely unknown. In this study, we generated, for the first time, nervous system-specific *Spns1* knockout mice, *Spns1^flox/flox^*;nestin-*Cre*, and found that the mutant mice develop neurological symptoms, such as epilepsy, and growth retardation, and die by 5 weeks of age. The mutant mice exhibited dysmyelination and oligodendrocyte shedding, while maintaining the neurons. Mutant mouse brains showed accumulation of lysophospholipids, predominantly in regions, such as the olfactory bulb and hippocampus. Furthermore, whereas sphingosine accumulated in the mutant mouse brain, the levels of ceramide and sphingoglycolipids, which are the main myelin components, were decreased. Our findings imply that abnormal sphingosine metabolism causes dysmyelination and white matter dysplasia in brain-specific *Spns1*-knockout mice, and indicate a possible role of SPNS1 mutation in the pathogenesis of congenital cerebral white matter dysplasia in humans.

## Introduction

Lysosomes, which are acidic organelles responsible for the degradation and recycling of various macromolecules taken up through endocytosis, phagocytosis, or autophagy, play a crucial role in cellular metabolism (Ballabio & Bonifacino, 2020; Perera & Zoncu, 2016). Proteins, polysaccharides, lipids, and nucleic acids are the substrates targeted for recycling in lysosomes. Lysosomes contain approximately 60 hydrolytic enzymes, including proteases, lipases, and nucleases (Bonam, Wang, & Muller, 2019). These hydrolytic enzymes are activated at an acidic pH of approximately 4.5, which is maintained via the activity of vacuolar H+-ATPase (v-ATPase), an ATP-driven proton pump located in the lysosomal membrane (Vasanthakumar & Rubinstein, 2020). Catabolites generated by lysosomal degradation are released into the cytosol through lysosomal transporters. These catabolites are subsequently used as new substrates in anabolic processes (Rudnik & Damme, 2021). Although several transporters for the export of amino acids, nucleosides, sugars, ions, and other moieties have been identified (Boswell-Casteel & Hays, 2017; Hu, Zhou, Cai, & Xu, 2022; Mayer et al., 2016; Meng, Heybrock, Neculai, & Saftig, 2020; van Veen et al., 2020; Wyant et al., 2017), lysosomal lipid transport remains poorly understood.

Sphingomyelin, ceramide, glycosphingolipids, and glycerophospholipids are recycled within lysosomes through the endocytosis pathway (Ogretmen, 2018). In this process, sphingolipids are enzymatically hydrolyzed into sphingosine, and glycerophospholipids, such as phosphatidylcholine (PC) and phosphatidylethanolamine (PE), are converted into lysophosphatidylcholine (LPC), lysophosphatidylethanolamine (LPE), and fatty acids (Thelen & Zoncu, 2017). Subsequently, sphingosine and lysoglycerophospholipids are released from the lysosomes. Several recent reports have revealed that protein spinster homolog 1 (SPNS1) transports LPCs from lysosomes in a proton gradient-dependent manner and re-esterifies them into PC in the cytosol, indicating that the phospholipid salvage pathway from the lysosomes to the cytosol is dependent on SPNS1 (Ha et al., 2024; He et al., 2022; Scharenberg et al., 2023). Physiologically, efflux of LPC by SPNS1 is required for cell survival under choline-limited conditions (Scharenberg et al., 2023). SPNS1 is also required for the release of sphingosine from lysosomes (Ha et al., 2024). Deletion of *Spns1* in the mouse liver results in lysosomal dysfunction accompanied by the accumulation of LPC and LPE in the lysosomes (He et al., 2022). *Spns1* systemic knockout mice become embryonically lethal between E12.5 and E13.5, and mice lacking *Spns1* postnatally show lipid accumulation in the lysosomes (Ha et al., 2024). Thus, there is growing evidence for the importance of SPNS1 in lysosomal homeostasis and for the association of SPNS1 abnormalities with lysosomal storage diseases (LSDs). Most LSDs follow a progressive neurodegenerative clinical course although symptoms in other organ systems are frequently encountered. However, the role of SPNS1 in central nervous system remains unclear.

In this study, we explored the role of SPNS1 in the central nervous system. We generated nervous system-specific *Spns1* knockout mice, *Spns1^flox/flox^*;nestin-*Cre* and found that these mice exhibited white matter dysplasia in the brain due to the loss of oligodendrocytes during the lactation period, resulting in death within 5 weeks of birth and severe growth retardation and epilepsy. Although the levels of LPC, LPE, lysophosphatidylinositol (LPI), and lysophosphatidylglycerol (LPG) were higher in mutant brains, those of galactosylceramide and sulfatide decreased compared with the levels in age-matched control brains, and fluctuations in their levels differed according to the acyl groups and brain regions. Swollen and dysfunctional lysosomes accumulated in SPNS1-deficient cells and tissues. Our data indicate that SPNS1 is indispensable for the metabolic pathway of specific lysophospholipids, whose loss makes oligodendrocytes vulnerable, and that SPNS1 is required for lysosomal function and integrity.

## Results

### Deletion of *SPNS1* results in loss of lysosomal integrity

To assess lysosomal integrity in the absence of *SPNS1*, we generated *SPNS1*-knockout HeLa cells (Fig. S1A). Immunofluorescence analysis using an antibody against LAMP-1, a lysosomal-associated membrane protein, showed that the number and size of lysosomes increased following the loss of *SPNS1* (Fig. 1A). Electron microscopy (EM) also showed the accumulation of enlarged lysosomes containing electron-dense structures in *SPNS1^-/-^* HeLa cells (Fig. 1B). The number of lysosomes was reduced following SPNS1-FLAG expression (Fig. 1A and B). Galectin-3, a beta-galactoside-binding lectin, which is a marker for lysosomal membrane rupture, was not detected in the lysosomes of either wild-type or *SPNS1^-/-^*HeLa cells under normal culture conditions (Fig. S1B). These results indicated that the loss of *SPNS1* is accompanied by persistent lysosomal dysfunction without membrane damage.

**Figure 1.**
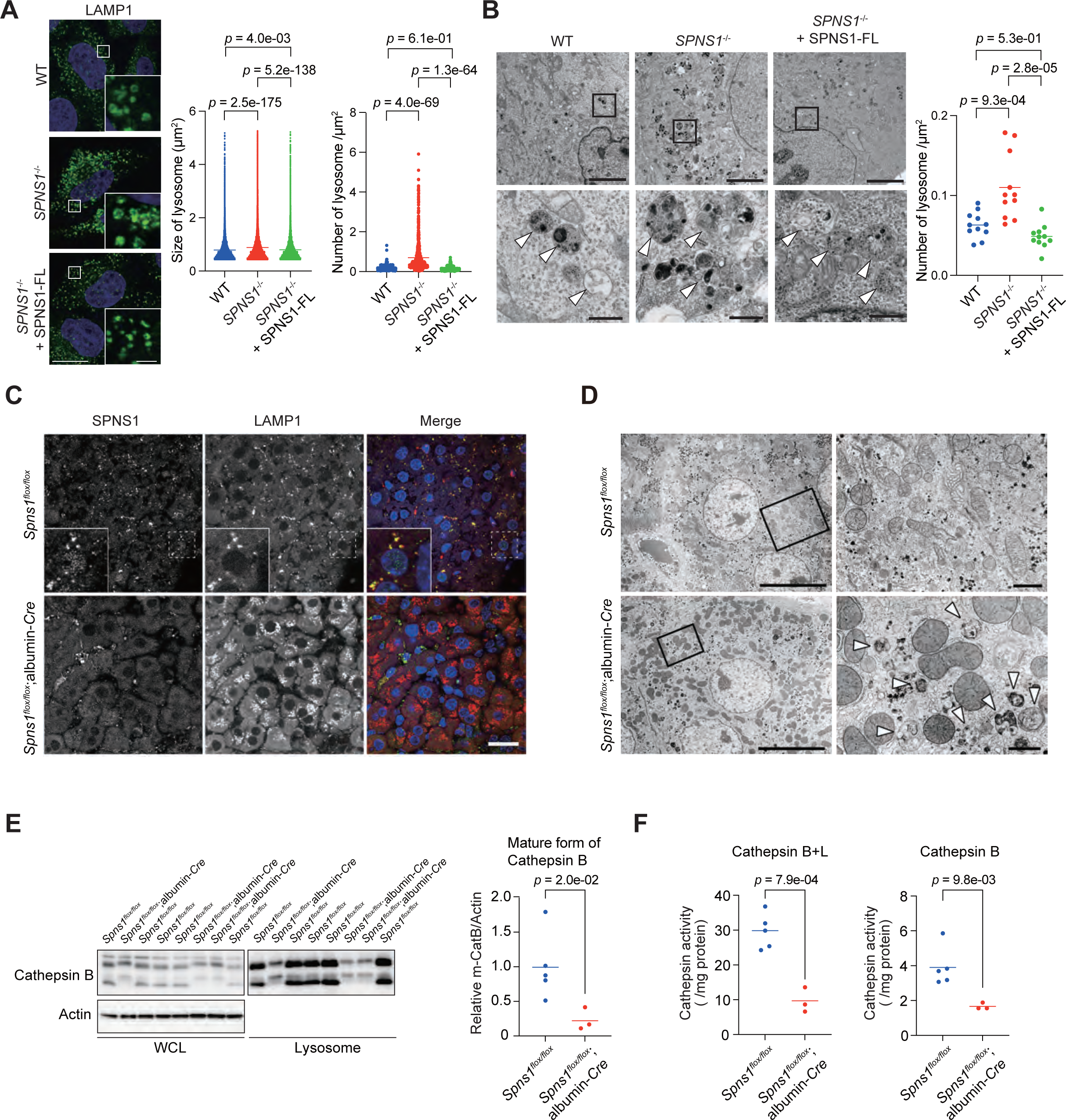
Impairment of lysosomal integrity by the loss of SPNS1. **(A)** Immunofluorescence analysis. Parental, *SPNS1^-/-^*HeLa cells and *SPNS1^-/-^* HeLa cells expressing SPNS1-FLAG were immunostained with the LAMP-1 antibody. The left scatter plot shows the results of quantitative analysis of the size of LAMP-1-positive structures per cell in parental (WT) (n = 34401), *SPNS1^-/-^* HeLa (n = 40011), and *SPNS1^-/-^*HeLa expressing SPNS1-FL (n = 34812). The right scatter plot shows the number of LAMP-1-positive structures per area (μm^2^) in WT (n = 663), *SPNS1^-/-^* HeLa (n = 740), and *SPNS1^-/-^* HeLa expressing SPNS1-FL (n = 463). Blue indicates nuclei stained with Hoechst 33342. Each dot corresponds to individual data points, and the horizontal line indicates the mean. Statistical analysis was performed using the Šidák’s test after one-way ANOVA. Scale bars, 20 μm (main panels), 2 μm (inset panels). **(B)** Electron micrographs of cells indicated in (A). Boxed regions are enlarged and shown below. The number of lysosomes per area (μm^2^) of WT (n = 11), *SPNS1^-/-^* HeLa (n = 11), and *SPNS1^-/-^*HeLa expressing SPNS1-FL (n = 11) were quantified using the Image J software and presented in scatter plots. Each dot corresponds to individual data points, and the horizontal line indicates the mean. Statistical analysis was performed using the Šidák’s test after one-way ANOVA. Arrowheads indicate lysosomes. Bars, top panels: 5 μm, bottom panels: 500 nm. **(C)** Immunofluorescence microscopy. Liver sections from *Spns1^flox/flox^* and *Spns1^flox/flox^*;albumin*-Cre* mice at 3 months were stained with SPNS1 (green) and LAMP-1 (red) antibodies, and nuclei was stained with Hoechst 33342 (blue) as indicated. Boxed regions are enlarged and shown in the insets. Bar, 20 μm. **(D)** Electron micrographs of mouse liver sections, as indicated. Boxed regions are enlarged and shown on the right. Arrows indicate lysosome-like structures with undigested materials. Bars, 10 μm (left) and 1 μm (right). **(E)** Immunoblot analysis. Homogenates and lysosomal fractions prepared from the liver of *Spns1^flox/flox^* (*n* = 5) and *Spns1^flox/flox^*;albumin*-Cre* mice (*n* = 3) at 6 weeks of age were subjected to SDS-PAGE followed by immunoblotting with Cathepsin B and Actin antibodies. Scatter plots show the results of densitometric analysis of the intensity of the Cathepsin B band relative to that of the Actin band. Each dot corresponds to individual data points, and the horizontal line represents the mean. Statistical analysis was performed using the Welch’s *t*-test. **(F)** Cathepsin activity. The activity of Cathepsin B + L and that of Cathepsin B in lysosomal fractions shown in (E) was measured. Each dot corresponds to individual data points, and the horizontal line represents the mean. Statistical analysis was performed using the Welch’s *t*-test.

To investigate the role of Spns1 in maintaining lysosomal integrity *in vivo*, we generated hepatocyte specific *Spns1*-knockout mice, *Spns1^flox/flox^*;albumin*-Cre*. Consistent with previous reports (Ha et al., 2024; He et al., 2022), the mutant mice exhibited mild liver injury, including slight but significant liver enlargement (Fig. S2A), and leakage of hepatic enzymes, which was observed continuously from 3 to 18 months of age. (Fig. S2B). Hematoxylin and eosin staining revealed slightly enlarged hepatocytes with mildly pale cytoplasm in the liver of 3-months-old *Spns1*-knockout mice. However, no significant abnormalities were observed in the tissue architecture (Fig. S2C). Immunostaining with the LAMP-1 antibody showed the accumulation of LAMP1-positive lysosomes in *Spns1*-knockout hepatocytes (Fig. 1C). Similar to *SPNS1*-deficient HeLa cells, hepatocytes from mutant mice contained many lysosome-like structures with undigested materials (Fig. 1D), suggesting impaired lysosomal digestion. The level of mature cathepsin B in lysosomal fractions of the liver from *Spns1*-knockout mice was much lower than that in the fractions from control mice liver (Fig. 1E), and the activities of both cathepsin B+L and cathepsin B in the liver from mutant mice were lower than those in the control (Fig. 1F). Taken together, these results indicate that SPNS1 is indispensable for lysosomal integrity.

### Loss of *Spns1* in the nervous system causes growth retardation and death in infancy

By crossbreeding *Spns1^flox/flox^* mice with transgenic mice expressing Cre recombinase under the control of nestin promoter (nestin*-Cre*), we generated nervous system-specific *Spns1*-deficient mice, *Spns1^flox/flox^*;nestin*-Cre*. Nestin promoter-driven Cre recombinase is expressed in the central and peripheral nervous systems, as well as in the precursors of neurons and glial cells. The Spns1 levels in the brain of *Spns1^flox/flox^*;nestin*-Cre* mice at postnatal day 12 (P12) were lower than that in age-matched brain of control mice (Fig. 2A). Mutant mice were born at Mendelian frequency and grew normally until 2 weeks of age; however, they showed obvious growth retardation (Fig. 2B) and often had epileptic seizures (Supplementary Movie). *Spns1^flox/flox^*;nestin*-Cre* mice showed decreased survival at 3 weeks of age, and all the mutant mice died by 5 weeks of age (Fig. 2C).

**Figure 2.**
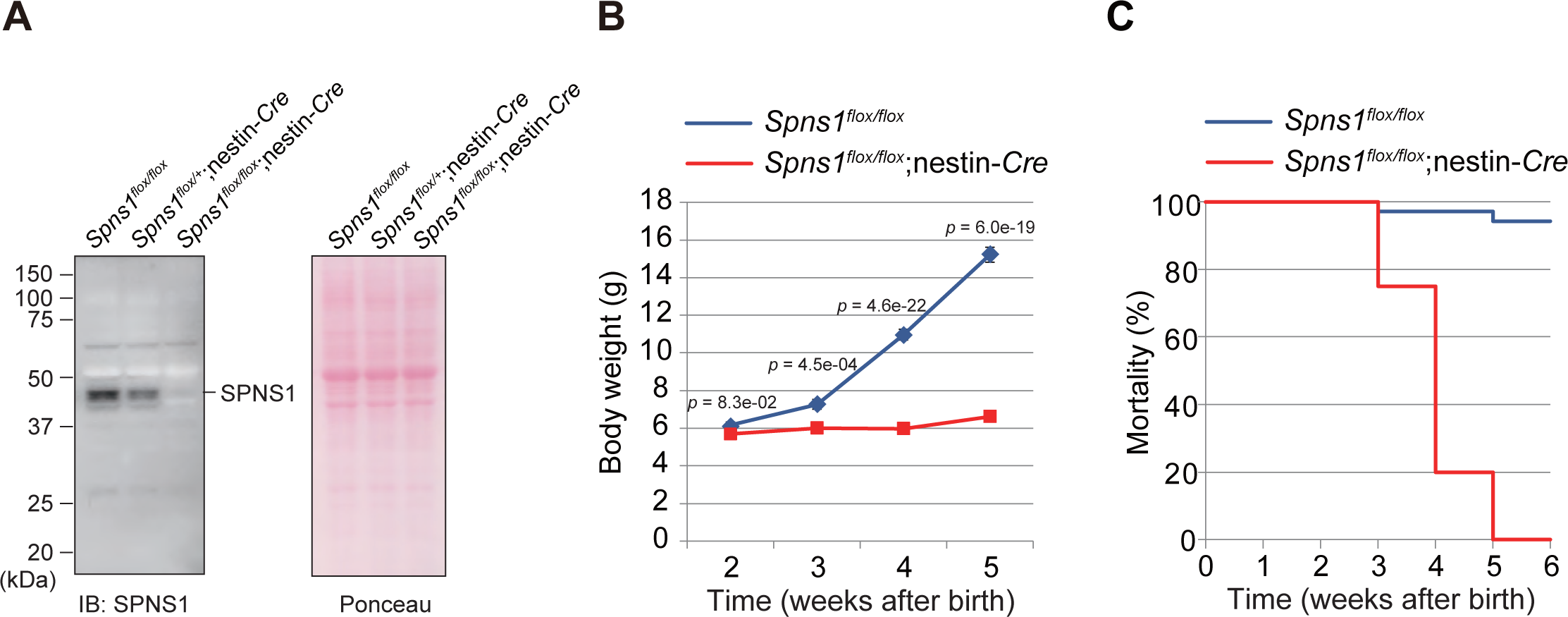
Growth and mortality of *Spns1^flox/flox^*;nestin*-Cre* mice. **(A)** Immunoblot analysis. Brain homogenates prepared from the indicated genotype mice were subjected to immunoblot analysis with the SPNS1 antibody and Ponceau-S staining of the gel after electrophoresis. Data shown are representative of three separate experiments. **(B)** Growth curve of *Spns1^flox/flox^* (*n* = 43 at 2 weeks of age, *n* = 41 at 3 weeks of age, *n* = 65 at 4 weeks of age, and *n* = 5 at 5 weeks of age) and *Spns1^flox/flox^*;nestin*-Cre* (*n* = 17 at 2 weeks of age, *n* = 18 at 3 weeks of age, *n* = 22 at 4 weeks of age, and *n* = 27 at 5 weeks of age) mice. Data are means ± s.e.m. Statistical analysis was performed using the Welch’s *t*-test. **(C)** Kaplan–Meier analysis of of *Spns1^flox/flox^* (*n* = 15) and *Spns1^flox/flox^*;nestin*-Cre* (*n* = 22) mice.

### Loss of *Spns1* in the nervous system causes dysmyelination due to the loss of oligodendrocytes

Next, we investigated the morphology of the brain in *Spns1^flox/flox^*;nestin*-Cre* mice. *Spns1*-deficient brain appeared smaller than control brain (Fig. 3A). Moreover, in the midsagittal section of brain from *Spns1^flox/flox^*;nestin*-Cre* mice, the white matter regions, especially the corpus callosum (Fig. 3A, arrowheads) and cerebellar medulla (Fig. 3A, arrows), were more transparent than those in the control brain (Fig. 3A). Hematoxylin and eosin (HE) staining revealed thinning of the corpus callosum in *Spns1^flox/flox^*;nestin*-Cre* mice (Fig. 3B), which suggested that the loss of Spns1 causes dysmyelination. Therefore, we performed immunofluorescence microscopy using an antibody against myelin basic protein (MBP), a major component of the myelin sheath. As shown in Fig. 3C, MBP signal intensity in the corpus callosum of *Spns1^flox/flox^*;nestin*-Cre* mice was almost completely lost. Consistent with this result, immunoblot analysis using the MBP antibody revealed that the amount of MBP in the brain of *Spns1^flox/flox^*;nestin*-Cre* mice was significantly lower than that in the control brain (Fig. 3D). Electron microscopy also showed fewer myelinated fibers in the corpus callosum of *Spns1^flox/flox^*;nestin*-Cre* mice compared with that in control mice (Fig. 3E). Given that oligodendrocytes are responsible for the formation of myelin sheath, these results suggest that Spin1 is involved in the differentiation and/or maturation of oligodendrocytes. To determine which developmental stages were affected, the number of oligodendrocytes was estimated during the lactation period by immunostaining with an antibody against OLIG2, a marker for immature and mature oligodendrocytes. At P0, the number of OLIG2-positive cells in the corpus callosum of *Spns1^flox/flox^*;nestin*-Cre* mice was comparable to that in control mice (Fig. 3F), suggesting that Spns1 is not involved in the differentiation of oligodendrocytes. However, at P18, the number was significantly lower than that in control mice (Fig. 3F). Although the decrease in the number of oligodendrocytes in Spns1-deficient mice was dramatic, we barely observed any abnormal tissue architecture or neuronal cell death in the mutant brain (Fig. S3A and B). We concluded that the loss of *Spns1* in the nervous system causes dysmyelination during the lactation period, which resuls in growth retardation and death in infancy.

**Figure 3.**
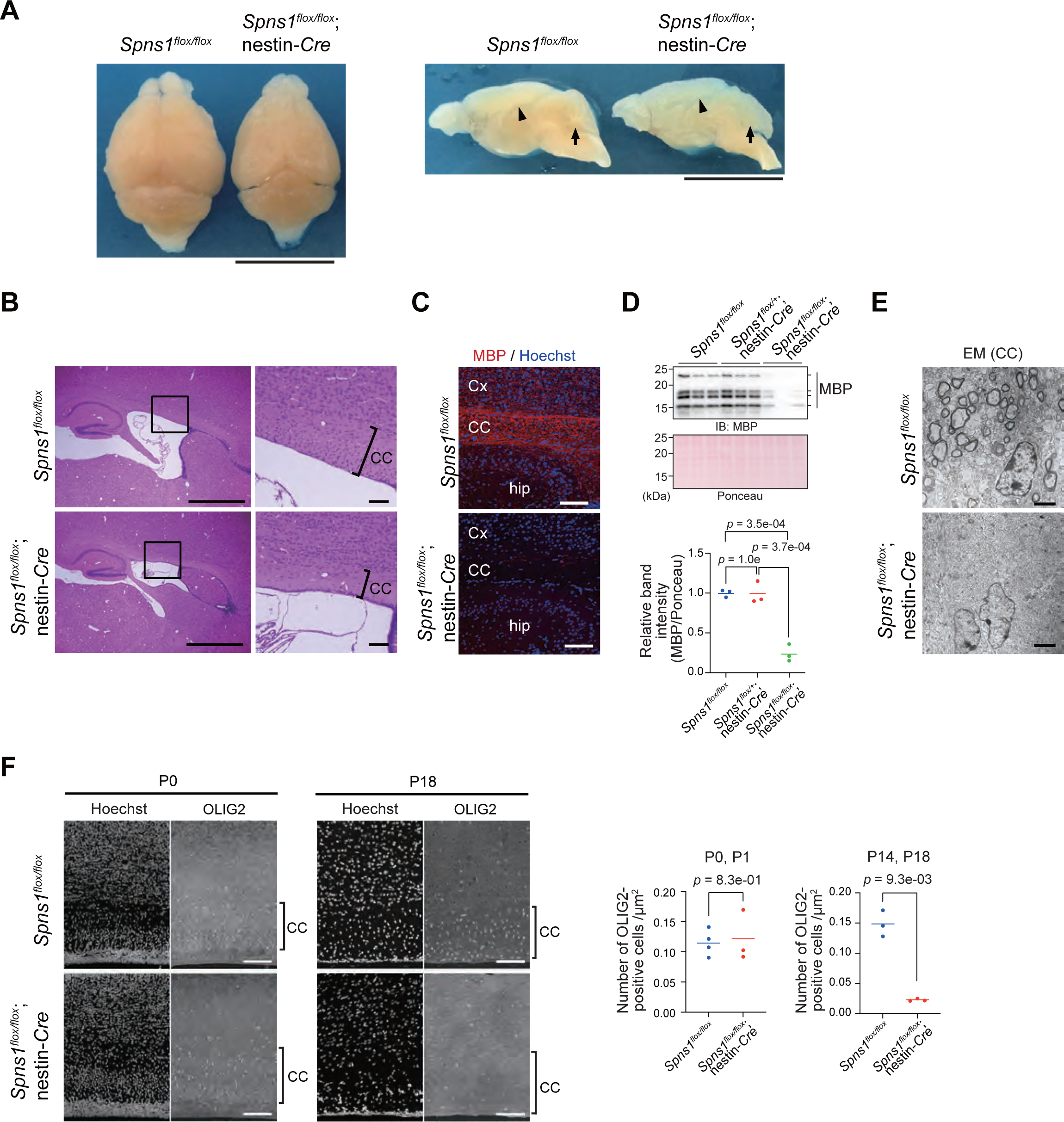
Dysmyelination due to the lack of oligodendrocytes in *Spns1^flox/flox^*;nestin*-Cre* mice. **(A)** Gross anatomy of whole brain (left) and sagittal sections (right) of brain from *Spns1^flox/flox^* and *Spns1^flox/flox^*;nestin*-Cre* mice at P26. Arrowheads indicate corpus callosum and arrows indicate cerebellar medulla. Bar, 1 cm. **(B)** Hematoxylin and eosin staining of the corpus callosum (CC) of brains from *Spns1^flox/flox^* and *Spns1^flox/flox^*;nestin*-Cre* mice at P24. Bars, 1 mm (left) and 100 μm (right). **(C)** Immunofluorescence images of sagittal sections of the brain from *Spns1^flox/flox^* and *Spns1^flox/flox^*;nestin*-Cre* mice at P26, stained with the MBP antibody (red) and Hoechst 33342 (blue). CC: corpus callosum; Cx: cerebral cortex; hip: hippocampus. Bar, 100 μm. **(D)** Immunoblot analysis. Homogenates of the brain tissue from *Spns1^flox/flox^*, *Spns1^flox/+^*;nestin*-Cre* and *Spns1^flox/flox^*;nestin*-Cre* mice at P12 were subjected to SDS-PAGE followed by immunoblotting with MBP and Ponceau-S staining of the gel. Scatter plots show the results of densitometric analysis of the MBP band relative to the whole protein content estimated using Ponceau-S staining (*n* = 3 each group). The dots represent individual data points, and the horizontal lines indicate the means. Statistical analysis was performed using the Šidák’s test after one-way ANOVA. **(E)** Electron micrographs of the corpus callosum (CC) of *Spns1^flox/flox^* and *Spns1^flox/flox^*;nestin*-Cre* mice at P14. Bar, 2 μm. **(F)** Immunofluorescence images of the sagittal sections of the brain from *Spns1^flox/flox^* and *Spns1^flox/flox^*;nestin*-Cre* mice at P0 (left) and P18 (right) stained with the OLIG2 antibody and Hoechst 33342. CC; corpus callosum. Bars, 100 μm. Scatter plots show the results of quantitative analysis of the number of OLIG2-positive cells. *Spns1^flox/flox^*(*n* = 4 at P0 and 1, *n* = 3 at P14 and 18, and *Spns1^flox/flox^*;nestin*-Cre* mice (*n* = 3 at P0 and 1, *n* = 3 at P14 and 18). The dots correspond to individual data points, and the horizontal lines represent the means. Statistical analysis was performed using the Welch’s *t*-test.

### Accumulation of diverse lysophospholipid molecular species in different regions of the brain of *Spns1*-deficient mice

Comprehensive lipidomic analysis was used to investigate the effect of Spns1 deficiency on the brain lipidome. An overview of the lipidome obtained using the hierarchical cluster analysis (HCA) revealed a massive accumulation of lysophospholipids, including LPCs, LPEs, and LPIs, in the brain of *Spns1^flox/flox^*;nestin*-Cre* mice (Fig. 4A). Volcano plot analysis also highlighted that loss of *Spns1* in the mouse brain resulted in the accumulation of lysophospholipids (Fig. 4B). We found that the levels of a wide variety of fatty acid-binding molecular species of several lysophospholipid molecular classes (LPCs, LPEs, and LPIs) were increased in the brain of *Spns1^flox/flox^*;nestin*-Cre* mice (Fig. 4C). Focusing on the major molecular species in each class, we observed that the accumulation was regardless of whether the fatty acids were saturated, monounsaturated, or polyunsaturated (Fig. 4C). Furthermore, imaging of LPCs showed that molecular species containing different fatty acids accumulated in different regions of the brain (Fig. 4D). Although all molecular species accumulated in parts of the olfactory bulb, arachidonoyl LPC, for example, showed significant accumulation in the cell layer of the hippocampus (Fig. 4D, arrow).

**Figure 4.**
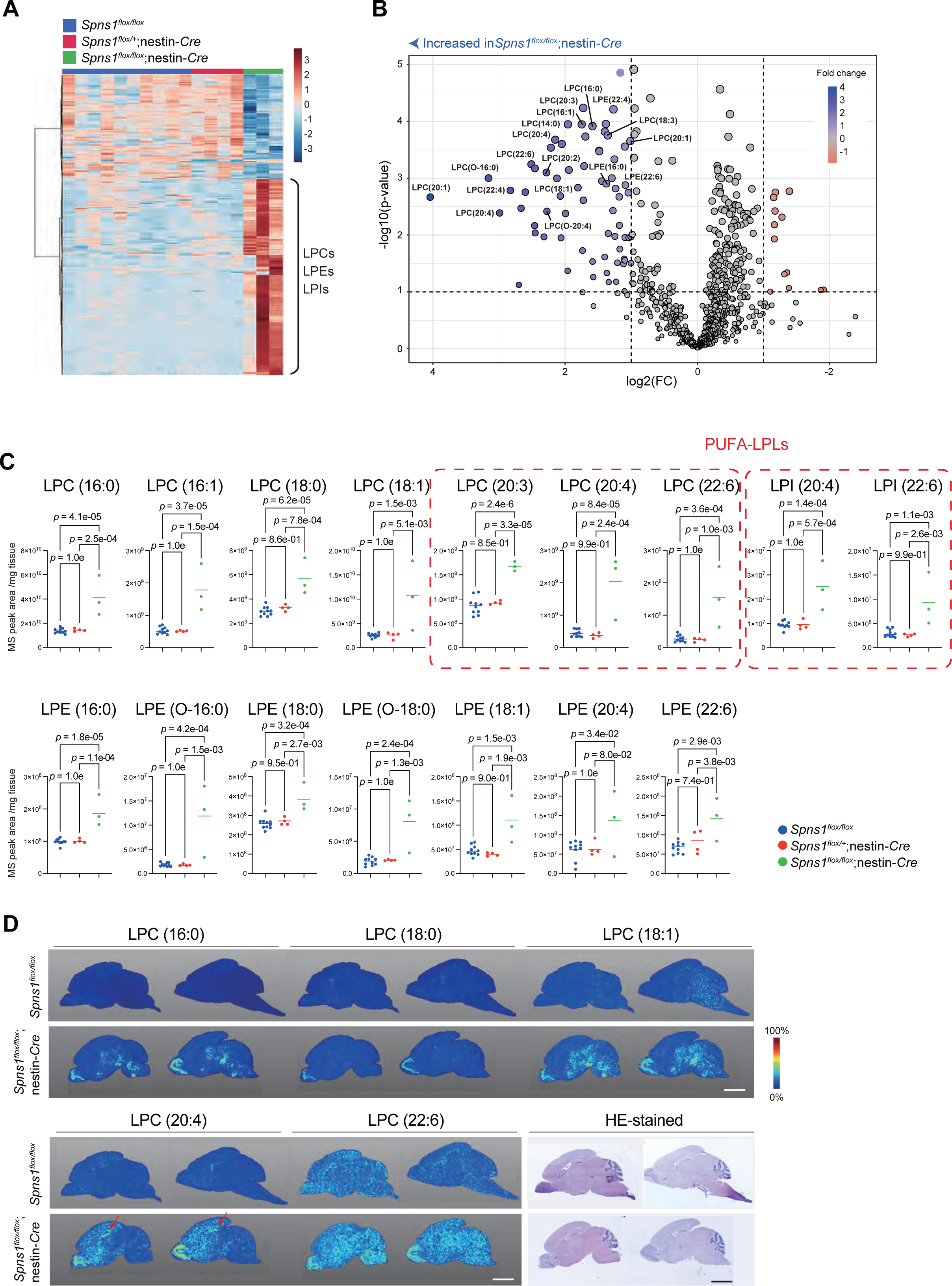
Quantification and visualization of lysophospholipids in *Spns1^flox/flox^*;nestin*-Cre* mice. **(A)** Lipidome analysis of the brain of *Spns1^flox/flox^* or *Spns1^flox/+^* (*n* = 10), *Spns1^flox/+^*;nestin*-Cre* (*n* = 4), and *Spns1^flox/flox^*;nestin*-Cre* (*n* = 3) mice at P14 was overviewed using hierarchical cluster analysis, which led to the identification of a cluster of accumulation of metabolites, including lysophosphatidylcholine (LPC), lysophosphatidylethanolamine (LPE), and lysophosphatidylinositol (LPI), in the brain of *Spns1^flox/flox^*;nestin*-Cre* mice. **(B)** Lipidome of the brain of *Spns1^flox/flox^* or *Spns1^flox/+^* (*n* = 10) and *Spns1^flox/flox^*;nestin*-Cre* (*n* = 3) mice at P14 was compared using the volcano plot analysis; among the molecular species with significant accumulation in *Spns1*-deficient brain, lysophospholipids were annotated. **(C)** Results of quantitative analysis of lysophospholipids, LPC, LPI, and LPE in the mouse brain described in (A); PUFA-containing LPs are highlighted with red dashed lines. The dots correspond to individual data points, and the horizontal lines represent the means. **(D)** Imaging mass spectrometry to visualize the pattern of lysophospholipid accumulation in sagittal sections of the brain of *Spns1^flox/flox^* and *Spns1^flox/flox^*;nestin*-Cre* mice at P14. Arrows indicate hippocampus. Bars, 3 mm.

### Accumulation of sphingosine and decrease in myelin glycolipid levels in the brain of *Spns1*-deficient mice

Besides lysophospholipids, accumulation of sphingosines and a corresponding decrease in the levels of ceramides, sphingomyelins, and sulfatides, which are synthesized from sphingosine, were observed in the brain of *Spns1^flox/flox^*;nestin*-Cre* mice (Fig. 5A). Because these sphingolipids constitute the myelin sheath formed by oligodendrocytes, we assessed the decrease in the levels of sphingolipids in the oligodendrocyte region (white matter) using imaging mass spectrometry. In the brain of *Spns1^flox/flox^*;nestin*-Cre* mice, a dramatic depletion of sphingolipids was noted in the white matter (Fig. 5B). Interestingly, sphingolipid depletion was more pronounced in the cerebral white matter than in the midbrain or hindbrain.

**Figure 5.**
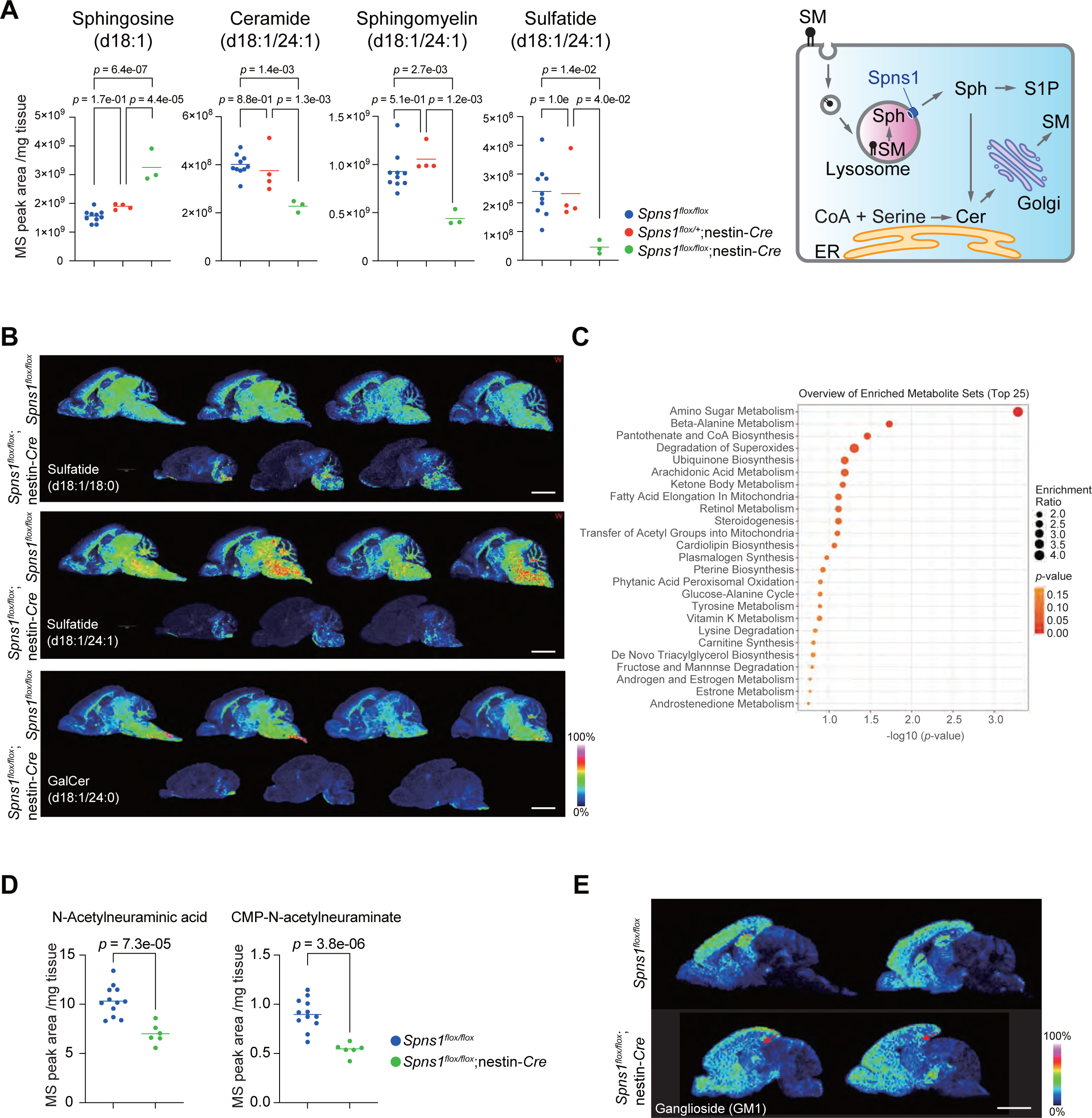
*Spns1*-deficient brain shows accumulation of sphingosine and decrease in myelin glycolipids. **(A)** Quantification of sphingosine and major sphingoglycolipids in the brain of *Spns1^flox/flox^* (*n* = 10), *Spns1^flox/+^*;nestin*-Cre* (*n* = 4), and *Spns1^flox/flox^*;nestin*-Cre* (*n* = 3) mice at P14. The dots correspond to individual data points, and the horizontal lines represent the means. A schematic illustration of sphingosine metabolic pathway. Sphingosine (Sph) are synthesized de novo synthesis in the Endoplasmic Reticulum (ER) or salvage pathway of the lysosome. Cer: ceramide; SM: sphingomyeline; S1P: sphingosine-1-phosphate; CoA: palmitoyl coenzyme A. **(B)** Imaging mass spectrometry to visualize the localization pattern of major sphingoglycolipid molecular species in the sagittal section of the brain of *Spns1^flox/flox^*and *Spns1^flox/flox^*;nestin*-Cre* mice at P14. Bars, 3 mm. **(C)** Results of the enrichment analysis of the dataset obtained from the water-soluble metabolome analysis of the brain comparing the brain of *Spns1^flox/flox^*and *Sns1^flox/flox^*;nestin*-Cre* mice at P14. **(D)** Quantification results of a sialic acid (N-acetyl neuraminic acid) and a corresponding sialic acid nucleotide (CMP-N-acetyl neuraminic acid) in the brain of *Spns1^flox/flox^*(*n* = 12) and *Spns1^flox/flox^*;nestin*-Cre* (*n* = 6) mice at P14. The dots correspond to individual data points, and the horizontal lines represent the means. **(E)** Imaging mass spectrometry to visualize the localization pattern of gangliosides in the sagittal sections of the brain of *Spns1^flox/flox^* and *Spns1^flox/flox^*;nestin*-Cre* mice at P14. Arrowheads indicate hippocampus. Bar, 3 mm.

In contrast, the hydrophilic brain metabolome showed abnormalities in the pathway responsible for glycosylation in *Spns1^flox/flox^*;nestin*-Cre* mice. As illustrated in Fig. 5C, an enrichment analysis overlooking the shift in the soluble metabolome showed that changes were enriched in the amino sugar metabolism pathway, which adds glycans to sphingolipids. In particular, the levels of sialic acid and sialic acid nucleotides in the brain of *Spns1^flox/flox^*;nestin*-Cre* mice were significantly lower than those in *Spns1^flox/flox^* mice (Fig. 5D). We then examined the localization of gangliosides, sialic acid-containing sphingolipids, using imaging mass spectrometry and observed a decrease in GM1 in the cerebrum, especially in the hippocampus (arrowheads in Fig. 5E).

## Discussion

Spns1 is a lysophospholipid transporter. However, because systemic *Spns1*-knockout mice are embryonically lethal (Ha et al., 2024), the role of Spns1 in the nervous system, particularly in the postnatal nervous system, is unknown. In this study, for the first time, we generated *Spns1^flox/flox^*;nestin*-Cre* mice, knocked out for *Spns1* in the nervous system. These mice showed neurological symptoms, such as epilepsy, and severe growth retardation and were dead within 5 weeks of birth (Figs. 2 and 3). Consistent with the results of lipidomic analysis of *Spns1*-deficient cultured cells, systemic *Spns1*-knockout embryos, and liver-specific *Spns1*-deficient mice (Ha et al., 2024; He et al., 2022; Scharenberg et al., 2023), our lipidomic analysis of the brain of nervous system-specific *Spns1*-knockout mice revealed accumulation of lysophospholipids, including LPCs, LPEs, and LPIs (Fig. 4). Using imaging MS, we found that lysophospholipids did not accumulate throughout the brain, but were abundant in limited regions, such as the olfactory bulb and hippocampus (Fig. 4). However, the effect of lysosomal accumulation of lysophospholipids in neurons may be limited because no neuronal cell death was detected in the regions of accumulation (Supplemental Fig. S3).

Spns1 also serves as a lysosomal transporter of sphingosines (Ha et al., 2024). Indeed, the brain of *Spns1*-knockout mice showed marked accumulation not only of lysophospholipids but also of sphingosine (Fig. 5). Moreover, the level of ceramide produced from sphingosine was decreased, as did the level of sphingoglycolipid synthesized from ceramide (Fig. 5). Sphingolipids are major components of the myelin sheath, and imaging MS revealed that they were prominent in the white matter of the corpus callosum and cerebellum in control mice, whereas no sphingolipid signals were obtained in the brain of mutant mice (Fig. 5). Considering that the mutant mice exhibited white matter dysplasia of the cerebrum and cerebellum (Fig. 3), it is likely that oligodendrocyte shedding occurred because of decreased sphingoglycolipid levels in these mice. Therefore, it is plausible that the defective transport of sphingosine, rather than of lysophospholipids, from lysosomes to the cytoplasm, followed by impairment of sphingoglycolipid biogenesis, is the primary cause underlying the dysmyelination phenotype of *Spns1^flox/flox^*;nestin*-Cre* mice.

In addition to the abnormalities in sphingoglycolipid metabolism, oligodendrocyte shedding, but not neuron shedding, was evident in *Spns1^flox/flox^*;nestin*-Cre* mice. Why does oligodendrocyte-specific shedding occur? During infancy, when large amounts of myelin proteins are synthesized, oligodendrocytes are subjected to ER stress (Clayton & Popko, 2016). Consistent with previous reports (Ha et al., 2024; He et al., 2022; Scharenberg et al., 2023), our analyses of *SPNS1*-deficient cultured cells and liver-specific *Spns1*-knockout mice showed lysosomal dysfunction (Fig. 1). In mutant oligodendrocytes, lysosomal dysfunction is superimposed on ER stress, which may be responsible for oligodendrocyte degeneration.

*Sialin*-deficient mice, which are incapable of transporting sialic acid from lysosomes, exhibit cytoplasmic sialic acid deficiency and insufficient ceramide synthesis that results in reduced sphingolipid levels and demyelination due to the loss of oligodendrocytes (Jhelum et al., 2020; Prolo, Vogel, & Reimer, 2009; Hu, 2023 #47; Traka, Podojil, McCarthy, Miller, & Popko, 2016) These mice are phenocopies of brain-specific *Spns1* knockout mice. Salla disease, a free sialic acid accumulation disorder, is an autosomal recessive genetic disorder caused by mutations in the SLC17A5 gene that encodes sialin. It is a neurodegenerative disease that results from the accumulation of free sialic acid within lysosomes (Ferreira & Gahl, 2017). Salla disease is usually asymptomatic at birth but develops in infancy. Developmental delay and growth retardation are present from early childhood, and approximately one-third of patients are able to walk independently. This condition is followed by a slow progression of ataxia, spasticity, athetosis, and degeneration of cognitive and motor functions into adulthood. Epilepsy may complicate the disease, and the absence of seizures is frequent. T2-weighted image shows extensive hyperintense white matter (hypomyelinating pattern), hypoplasia of the corpus callosum, and atrophy of the cerebellum, suggesting abnormal myelination of the cerebrum (Gupta et al., 2023). Notably, only approximately two-thirds of patients clinically diagnosed with congenital cerebral white matter dysplasia, including those diagnosed with Salla disease, have causative gene mutations, which indicates that other unidentified disease-causing genes might be involved. Recently, a homozygous hypomorphic variant of *SPNS1* was identified in three patients with developmental delay, neurological symptoms, intellectual disability, and cerebellar hypoplasia (Ha et al., 2024). Our data suggest that *SPNS1* is a causative gene for congenital cerebral white matter dysplasia and that the pathogenesis of Salla disease and *SPNS1* mutations in humans is due to abnormal sphingosine metabolism.

Limitations of this study are that it is unclear whether the shedding of *Spns1*-deficient oligodendrocytes *in vivo* is truly caused by a combination of ER and lysosomal stresses, and that the *SPNS1* mutation has not been proven to be the causative gene for Salla disease.

## Materials and methods

### Cell culture

HeLa cells (ATCC CCL2} were grown in Dulbecco’s modified Eagle medium (DMEM) supplemented with 10% fetal bovine serum, 2 mM L-glutamine, 5 U/mL penicillin, and 50 μg/mL streptomycin. *SPNS1* (5′-gacgacgggccagtgcctgg-3′) guide RNA was designed using the CRISPR Design tool (http://crispr.mit.edu/) and subcloned into pX330-U6-Chimeric_BB-CBh-hSpCas9 (Addgene #42230), a human codon-optimized SpCas9 and chimeric guide RNA expression plasmid. To generate *SPNS1* knockout HeLa cells, the cells were transfected with the aforementioned pX330 vectors together with pEGFP-C1 (#6084-1, Clontech Laboratories, Mountain View, CA, USA) and cultured for 2 d. Green fluorescent protein (GFP)-positive cells were sorted and expanded. Ablation of *SPNS1* was confirmed using a heteroduplex mobility assay, followed by immunoblot analysis with an SPNS1 antibody. HeLa cells were authenticated using the STR profile. All the cell lines were tested for mycoplasma contamination.

### Mice

*Spns1^flox/flox^* mice were generated on a C57BL/6 × CBA background. Two *loxP* sites flanking exons 1 and 2 of *Spns1* were introduced and a neomycin-resistance cassette (neo) flanked by *FRT* sites was inserted between exon 2 and the second *loxP* site to prepare the targeting vector. The linearized targeting construct was electroporated into TT2 embryonic stem cells to obtain the recombinant Neo allele. Germline transmission of the Neo allele was confirmed using PCR and Southern blot analyses. Mice with the floxed *Spns1* allele were generated by mating mice with the Neo allele with *FLPe* transgenic mice. The resulting *Spns1^flox/flox^* mice were crossed with nestin-*Cre* and albumin-*Cre* mice to generate nervous system- and hepatocyte-specific *Spns1* knockout mice, respectively. The mice were housed in specific pathogen-free facilities. The experimental protocols were approved by the Ethics Review Committee for Animal Experimentation of Juntendo University (2024240, 2024241; approved March 18, 2024).

### Immunoblot analysis

Mouse brain and liver were homogenized in 0.25 M sucrose, 10 mM 2-[4-(2-hydroxyethyl)-1-piperazinyl]ethanesulfonic acid (HEPES) (pH 7.4), and 1 mM dithiothreitol (DTT). HeLa cells were lysed in ice-cold TNE buffer (50 mM Tris-HCl [pH 7.5], 150 mM NaCl, and 1 mM EDTA) containing 1% NP40, 1% Triton X-100, and protease inhibitors. The lysates were centrifuged at 20,000 × *g* for 10 min at 4°C, and the resulting supernatants were used as samples for immunoblot analysis. Samples were subjected to SDS-PAGE and transferred onto a polyvinylidene difluoride membrane (IPVH00010; Merck Millipore, Burlington, MA, USA). Antibodies against SPNS1 (HPA041995; Human Protein Atlas, Bromma, Sweden; 1:500), MBP (MCA409S; Bio-Rad, Hercules, CA, USA; 1:500), TFEB (4240; Cell Signaling Technology, Danvers, MA, USA; 1:500), Cathepsin B (219361; Calbiochem, San Diego, CA, USA; 1:500), and ACTIN (A1978; Sigma-Aldrich, Burlington, MO, USA; 1:2000) were purchased from the indicated suppliers. The blots were incubated with a horseradish peroxidase-conjugated goat anti-mouse IgG (H+L) antibody (115-035-166, Jackson ImmunoResearch Laboratories, Inc. West Grove, PA, USA; 1:10000) or goat anti-rabbit IgG (H+L) (111-035-144, Jackson ImmunoResearch Laboratories, Inc.; 1:10000), and visualized using chemiluminescence. Band density was measured using the Multi Gauge V3.2 software (FUJIFILM Wako Pure Chemical Corporation, Osaka, Japan).

### Immunofluorescence analysis

HeLa cells cultured on coverslips were washed with phosphate-buffered saline (PBS), fixed with 4% paraformaldehyde (PFA) for 15 min at 24°C, permeabilized with 0.1% digitonin in PBS for 5 min, and blocked with TNB blocking buffer (50 mM Tris-HCl [pH 7.5], 150 mM NaCl, and 0.5% TSA blocking reagent [FP102, PerkinElmer, MA, USA]) for 30 min. The cells were then incubated with primary antibodies in the blocking buffer for 1 h, washed with PBS, and incubated with secondary antibodies for 1 h. Antibodies against TFEB (4240, Cell Signaling Technology 1:200), LAMP-1 (AF4800, R&D Systems, Minneapolis, MN, USA; 1:200), and GALECTIN-3 (sc-23938, Santa Cruz Biotechnology Inc.; 1:200) were used as primary antibodies. Goat anti-mouse IgG (H + L) Highly Cross-Adsorbed Secondary Antibody, Alexa Fluor 647 (A21236, Thermo Fisher Scientific) and goat anti-rabbit IgG (H + L) Cross-Adsorbed Secondary Antibody, Alexa Fluor 488 (A11008, Thermo Fisher Scientific) were used as secondary antibodies. The cells were imaged using an FV3000 confocal laser-scanning microscope with FV31S-SW (version: 2.4.1.198) (Olympus, Tokyo, Japan) and a UPlanXApo ×60 NA 1.42 oil objective lens. The contrast and brightness of the images were adjusted using Photoshop 2021v25.0 (Adobe, San Jose, CA, USA). The number and size of LAMP-1-positive punctae in each cell and the mean fluorescence intensity of TFEB-positive punctae were quantified using a Benchtop High-Content Analysis System (CQ1; Yokogawa Electric Corp., Tokyo, Japan) and the CellPathfinder software (Yokogawa Electric Corp.).

### Measurement of cathepsin activity

Lysosome fractions were prepared from the liver of *Spns1^flox/flox^* and *Spns1^flox/flox^*;albumin*-Cre* mice in accordance with a previously published method with minor modifications (Ueno, Muno, & Kominami, 1991). Briefly, the liver was excised and mashed through a stainless-steel mesh. The mashed tissue was suspended in four volumes of ice-cold 5 mM N-tris(hydroxymethyl)methyl-2-aminoethanesulfonic acid (TES) (pH 7.5) containing 0.3 M sucrose (TES buffer) and homogenized using a Dounce homogenizer on ice. The homogenate was centrifuged at 700 × *g* for 5 min at 4°C, and the postnuclear supernatant was carefully collected. The pellet was suspended with 5 mL of TES buffer, and the homogenate was recentrifuged at 700 × *g* for 5 min at 4°C. The resulting supernatant was combined with the postnuclear supernatant and centrifuged at 12,000 × *g* for 5 min at 4°C. The pelleted mitochondrial-lysosomal fraction was suspended in 500 μL of TES buffer. This suspension was loaded onto 25 mL of 57% Percoll containing TES buffer and centrifuged at 50,000 × *g* for 45 min. After centrifugation, 4 mL fractions were collected from the bottom of the tube, diluted with 40 mL of TES buffer, and centrifuged at 10,000 × *g* for 20 min at 4°C. The resulting pellet was suspended in 700 μL of TES buffer. The cathepsin activity of the lysosomal fraction was measured fluorometrically by hydrolysis of z-Phe-Arg-MCA (Peptide Institute, 3095-v) to quantify cathepsin B and L, and that of z-Arg-Arg-MCA (Peptide Institute, 3123-v) to quantify cathepsin B activity. Briefly, the lysosomal fraction was suspended in TES buffer and incubated with the substrates at 37°C for 10 min. The reaction was stopped by the addition of 5% SDS and 0.1 M Tris-HCl (pH 9.0), and the fluorescence emission was measured at 470 nm after excitation at 370 nm.

### Histological analyses

The mouse brain was fixed by perfusion with 4% PFA–4% sucrose in 0.1 M phosphate buffer (PB) (pH 7.4) and embedded in paraffin. Three micrometer-thick paraffin sections were prepared and processed for HE staining or immunohistochemical fluorescence microscopy. For immunostaining, antigen retrieval was performed for 20 min at 98°C using a microwave processor (MI-77; AZUMAYA, Tokyo, Japan) in 1% immunosaver (Nissin EM, Tokyo, Japan). The sections were blocked and incubated for 2 days at 4°C with the following primary antibodies: mouse anti-MBP (sc-66064, Santa Cruz Biotechnology Inc.), goat anti-OLIG2 (Bio-Techne, MN, USA), rabbit anti-SPNS1 (HPA041995, Sigma-Aldrich, MO, USA), and anti-rat Lamp1 (sc-19992, Santa Cruz Biotechnology Inc., sc-19992). The sections were further incubated with Alexa Fluor 594- or Alexa Fluor 488-conjugated donkey anti-mouse IgG, donkey anti-goat IgG, and/or donkey anti-rat IgG (Jackson ImmunoResearch Laboratories). Immunofluorescence images were obtained with a laser scanning confocal microscope (FV1000; Olympus) equipped with a 20× (UPlanSApo 20×, NA 0.75) or a 60× (PlanApo N 60×, NA 1.42 oil) objective lens, or a microscope (BX51, Olympus, Japan) equipped with a cooled CCD camera system (DP-71, Olympus) and a 4× (UPlanSApo 4×, NA 0.16) or a 10× (UPlanSApo 10×, NA 0.40) objective lens. After image acquisition, the contrast and brightness were adjusted using Photoshop CS6 (Adobe).

### Electron microscopy (EM)

For conventional EM, mouse brain was fixed by perfusion with 2% PFA–2% glutaraldehyde in 0.1 M PB, pH 7.4, and portions of the corpus callosum were excised. HeLa cells were fixed using the same fixative. They were further processed in accordance with the reduced-osmium method and embedded in Epon812. Ultrathin sections were prepared, stained with uranyl acetate and lead citrate, and observed under an electron microscope (JEM1400EX, JEOL).

### Lipidome analysis

Freshly excised brain tissue was immediately snap-frozen and preserved at −80°C until further processing. For total lipid extraction (Alshehry, 2015), the brain tissue was homogenized in 1000 μL of a 1:1 (v/v) 1-butanol/methanol solution containing 5 mM ammonium formate with a manual homogenizer (Finger Masher, AM79330, Sarstedt, Tokyo, Japan). After homogenization for a few minutes, the homogenate was centrifuged at 16,000 × *g* for 10 min at 20°C and the supernatant was collected and transferred into a 200 μL LC-MS vial. For lipidome analysis, an Orbitrap-based MS (Q-Exactive Focus, Thermo Fisher Scientific, San Jose, CA, USA) coupled to an HPLC system (Ultimate3000, Thermo Fisher Scientific) was used. The chromatographic and mass spectrometric parameters were adapted from a previously described method (Ruzicka, 2014). Elution was performed on a Thermo Scientific Accucore C18 column (2.1 × 150 mm, 2.6 μm). The mobile phase consisted of phase A (10 mM ammonium formate in 50% acetonitrile (v/v) and 0.1% formic acid (v/v)) and phase B (2 mM ammonium formate in a mixture of acetonitrile:isopropyl alcohol:water at a ratio of 10:88:2 (v/v/v) with 0.02% formic acid). The elution gradient (phase A:phase B) was set as follows: 65:35 at 0 min, 40:60 from 0 to 4 min, 15:85 from 4 to 12 min, 0:100 from 12 to 21 min, maintained at 0:100 from 21 to 24 min, changed to 65:35 at 24.1 min, and finally 100:0 from 24.1 to 28 min. The flow rate was 0.4 mL/min and the column temperature was maintained at 35°C.

The Q-Exactive Focus Mass Spectrometer was operated in positive and negative ESI modes. A full mass scan (*m/z* 250–1100), followed by data-dependent MS/MS, was performed at resolutions of 70,000 and 17,500. The automatic gain control target was set at 1 × 10^6^ ions, and maximum ion injection time was 100 ms. Source ionization parameters were as follows: spray voltage, 3 kV; transfer tube temperature, 285°C; S-Lens level, 45; heater temperature, 370°C; sheath gas, 60; and auxilliary gas, 20. Acquired data were analyzed using the LipidSearch software (Mitsui Knowledge Industry, Tokyo, Japan) with the following parameters: precursor mass tolerance, 3 ppm; product mass tolerance, 7 ppm; and m-score threshold, 3.

### Extraction of metabolites from the brain tissue for metabolomic analysis

For detailed metabolomic analysis, metabolites from the brain tissue were extracted using a previously described method with slight modifications (Maeda et al., 2023). Frozen tissue samples were homogenized in 500 μL of ice-cold methanol containing methionine sulfone (L-Met) and 2-morpholinoethanesulfonic acid (MES) as internal standards for cationic and anionic metabolites, respectively, using a Finger Masher manual homogenizer (AM79330; Sarstedt, Tokyo, Japan). The homogenate was then mixed with half the sample volume of ultrapure water (LC/MS grade, sourced from Wako) and 0.4-times the original volume of chloroform (Nacalai Tesque, Kyoto, Japan). The mixture was centrifuged at 15,000 × *g* for 90 min at 4°C. After centrifugation, the aqueous layer was collected and filtered through an ultrafiltration tube (UltraFree MC-PLHCC; Human Metabolome Technologies, Yamagata, Japan). The filtered aqueous extract was evaporated under a stream of nitrogen using a heating block (DTU-28N; TAITEC, Koshigaya City, Japan). Finally, the concentrated sample was reconstituted in 50 μL of ultrapure water for further analysis using ion chromatography-high resolution mass spectrometry (IC-HR-MS).

### Metabolome analysis using IC-HR-MS

For metabolite detection, we used an Orbitrap-type MS (Q-Exactive focus; Thermo Fisher Scientific) coupled with a high-performance IC system (ICS-5000+, Thermo Fisher Scientific). The IC unit was equipped with an anion electrolytic suppressor (Dionex AERS 500; Thermo Fisher Scientific) that converted the potassium hydroxide gradient to pure water prior to the introduction of sample into the MS system. Metabolite separation was performed using a Dionex IonPac AS11-HC column with 4 μm particle size (Thermo Fisher Scientific). The flow rate for IC was set at 0.25 mL/min, and it was supplemented post-column with a 0.18 mL/min makeup flow of methanol. The potassium hydroxide gradient for IC was adjusted as follows: start at 1 mM, increased to 100 mM over 40 min, maintained at 100 mM for the next 10 min, and then returned to 1 mM over the final 10 min. The column temperature was maintained at 30°C. The mass spectrometer was operated in ESI-positive and ESI-negative modes for all measurements, and a full mass scan from m/z 70 to 900 was conducted at a resolution of 70,000. The automatic gain control was set at a target of 3 × 10^6^ ions, with a maximum ion injection time of 100 ms. The optimized ionization parameters were as follows: spray voltage, 3 kV; transfer temperature, 320°C; S-Lens level, 50; heater temperature, 300°C; sheath gas flow, 36; and auxiliary gas flow, 10.

### Multivariate statistical analysis

To evaluate the differences in the serum metabolome between the experimental groups and identify significant metabolite contributors, HCA and volcano plot analysis were performed using Metaboanalyst (v4.0), an online tool for multivariate statistical analyses. For normalization, the samples were subjected to median adjustment to uniformly correct for systematic differences across the samples. Autoscaling was applied to standardize the comparisons of variables, without performing data transformation.

### Sample preparation for imaging mass spectrometry (IMS)

For IMS, the brain tissue was embedded in super cryoembedding medium (SCEM; SECTION LAB, Hiroshima, Japan) and subsequently stored at −80°C. Tissue blocks containing SCEM were sectioned at −16°C into 8-micrometer slices using a CM 3050 cryostat (Leica, Wetzlar, Germany). The slices were then transferred onto indium tin oxide-coated glass slides (Matsunami Glass Industries, Osaka, Japan) for subsequent processing. The sections were manually coated with a matrix solution of 2,5-dihydroxy benzoic acid (50 mg/mL in 80% ethanol) and 9-aminoacridine (10 mg/mL in 80% ethanol) for positive and negative ion detection, respectively, using an art brush (Procon Boy FWA Platinum; Mr. Hobby, Tokyo, Japan). The matrix was sprayed onto the slides from a distance of approximately 15 cm, applying approximately 1 mL of the solution per slide. To ensure consistent conditions for analyte extraction and cocrystallization, the matrix was simultaneously applied across multiple slides. Optical images of the sections were captured using a scanner and analyzed using matrix-assisted laser desorption/ionization (MALDI)-MS imaging.

### IMS

MALDI imaging was performed using a Bruker timsTOF fleX MS (Bruker Daltonics, Bremen, Germany) operating in the quadrupole time-of-flight (qTOF) mode. The setup parameters included detection in positive and negative ion modes, a pixel resolution set at 80 μm, 200 laser pulses per pixel at a frequency of 10 kHz, and the laser power adjusted to 50%. The data collection focused on an m/z range of 100–650. The obtained raw mass spectra were processed and visualized using the SCiLS Lab software (v. 2019, Bruker Daltonics), enabling the production of detailed MS images. The signals within the specified m/z range were normalized to the total ion current to mitigate variations in ionization efficiency across different pixels. Metabolite identities were verified through accurate mass measurements and by comparison with reference standards MALDI-MS.

### Statistical analysis

Statistical analyses were performed using the unpaired *t*-test (Welch’s *t*-test), Šidák’s multiple comparison test after one-way ANOVA. GraphPad PRISM 10 (GraphPad Software) was used for statistical analyses. All tests were two-sided, and *P*-values <0.05 were considered statistically significant.

## Acknowledgements

Funding statement: This work was supported by JSPS KAKENHI Grant Numbers JP19H05706, JP21H004771, 23K20044, 24H00060 (to M.K.), 23K06324 (to Y-s.S.), 23K06415 (to Y.I.), 17K07380 (to Y.K.), JP20H03415 (to S.W.), JP23H02660 (to S.W.); AMED Grant Number JP22gm1410004h0003 (to M.K.); the Takeda Science Foundation (to M.K.); the Uehara Memorial Foundation (to M.K.); and by the Kobayashi Foundation (to M.K.). This work was also supported by JSPS KAKENHI Grant Number JP22H04926, Grant-in-Aid for Transformative Research Areas ― Platforms for Advanced Technologies and Research Resources “Advanced Bioimaging Support.” We thank H. Morishita, J. Sakamaki, and K. Tabata for critically reading this manuscript. We also thank A. Yabashi and K. Kanno for their help with the morphological analyses. We would like to thank Editage (www.editage.jp) for English language editing.

Conflicts of Interest: We declare that we have no competing financial interests. Author Contributions: M.K. designed and directed the study. Y.I., Y.K., and Y-s.S. performed the biochemical experiments. Y.S. performed the lipidome and mass spectrometry imaging. Y-s.S., T.U., N.T., and M. Koike, S.W. performed the EM and histological analyses. S.K-H. generated KO cell lines. All the authors discussed the results and commented on the manuscript. Data availability: All data required to evaluate the conclusions of this study are presented in the paper and/or Supplementary Materials.

---

**Figure S1. Lysosomal membrane integrity of *SPNS1*-knockout HeLa cells.**

**(A)** Generation of *SPNS1*-knockout HeLa cells. Parental and *SPNS1*-deficient HeLa cells were lysed, and then subjected to SDS-PAGE followed by immunoblot analysis with the SPNS1 antibody. Data are representative of three separate experiments. **(B)** Immunofluorescence analysis. Parental (WT), *SPNS1^-/-^* HeLa cells and *SPNS1^-/-^* HeLa cells expressing SPNS1-FLAG were immunostained with GALECTIN-3 and LAMP-1 antibodies under normal and L-leucyl– L-leucine methyl ester (LLOMe)-treated conditions. LLOMe is a lysosomotropic agent that severely damages and permeabilizes the lysosomal membrane. Lysosomes in both control and *SPNS1*-knockout HeLa cells were positive for GALECTIN-3 upon LLOMe treatment. Blue indicated nuclei stained with Hoechst 33342. Bars, 20 μm and 2 μm.

**Figure S2. Phenotype of hepatocyte-specific *Spns1*-knockout mice.**

**(A)** Liver weight (% per body weight) in *Spns1^flox/flox^*(3-month-old, *n* = 4; 9-month-old, *n* = 16; 12-month-old, *n* = 14; 18-month-old, *n* = 10) and *Spns1^flox/flox^*;albumin-*Cre* (3-month-old, *n* = 3; 9-month-old, *n* = 4; 12-month-old, *n* = 17; 18-month-old, *n* = 7) mice. The dots correspond to individual data points, and the horizontal lines represent the means. Statistical analysis was performed using the Šidák’s test after one-way ANOVA. **(B)** Liver function tests in mice described in (A). The serum levels of aspartate aminotransferase (AST), alanine aminotransferase (ALT), and alkaline phosphatase (ALP) were measured. IU/L, international units/liter. The dots represent individual data points, and the horizontal lines indicate the means. Statistical analysis was performed using the Šidák’s test after one-way ANOVA. **(C)** Hematoxylin and eosin staining of liver paraffin sections from 3-month-old *Spns1^flox/flox^*and *Spns1^flox/flox^*;albumin-*Cre* mice. Boxed regions are enlarged and shown in insets. C: central vein; P: portal vein. Bars: 50 µm and 20 µm (inset).

**Figure S3.** Few abnormalities in *Spns1*-deficient neurons.

Hematoxylin and eosin staining of the olfactory bulb **(A)** and hippocampal formation **(B)** from 2-week-old *Spns1^flox/flox^*and *Spns1^flox/flox^*;nestin-*Cre* mice. The boxed regions are enlarged and shown below, as indicated. M, mitral cell layer; DG, dentate gyrus. Bar, 200 μm (low magnification) and 20 μm (high magnification).

## Notes

### Competing Interest Statement

The authors have declared no competing interest.

## References

Alshehry, Z. H. (2015). An efficient single phase method for the extraction of plasma lipids. Metabolites, 5. doi:10.3390/metabo5020389

Ballabio, A., & Bonifacino, J. S. (2020). Lysosomes as dynamic regulators of cell and organismal homeostasis. Nat Rev Mol Cell Biol, 21(2), 101–118. doi:10.1038/s41580-019-0185-4

Bonam, S. R., Wang, F., & Muller, S. (2019). Lysosomes as a therapeutic target. Nat Rev Drug Discov, 18(12), 923–948. doi:10.1038/s41573-019-0036-1

Boswell-Casteel, R. C., & Hays, F. A. (2017). Equilibrative nucleoside transporters-A review. Nucleosides Nucleotides Nucleic Acids, 36(1), 7–30. doi:10.1080/15257770.2016.1210805

Clayton, B. L. L., & Popko, B. (2016). Endoplasmic reticulum stress and the unfolded protein response in disorders of myelinating glia. Brain Res, 1648(Pt B), 594–602. doi:10.1016/j.brainres.2016.03.046

Ferreira, C. R., & Gahl, W. A. (2017). Lysosomal storage diseases. Transl Sci Rare Dis, 2(1-2), 1–71. doi:10.3233/TRD-160005

Gupta, J., Reddy, N., Mankad, K., Kabra, U., Bhandari, A., Bagarhatta, M., & Gupta, A. (2023). Teaching NeuroImage: A New Imaging Finding in a Boy With Salla Disease Caused by a Pathogenic Variant in the SLC17A5 Gene. Neurology, 101(14), 631–632. doi:10.1212/WNL.0000000000207546

Ha, H. T., Liu, S., Nguyen, X. T., Vo, L. K., Leong, N. C., Nguyen, D. T., … Nguyen, L. N. (2024). Lack of SPNS1 results in accumulation of lysolipids and lysosomal storage disease in mouse models. JCI Insight, 9(8). doi:10.1172/jci.insight.175462

He, M., Kuk, A. C. Y., Ding, M., Chin, C. F., Galam, D. L. A., Nah, J. M., … Silver, D. L. (2022). Spns1 is a lysophospholipid transporter mediating lysosomal phospholipid salvage. Proc Natl Acad Sci U S A, 119(40), e2210353119. doi:10.1073/pnas.2210353119

Hu, M., Zhou, N., Cai, W., & Xu, H. (2022). Lysosomal solute and water transport. J Cell Biol, 221(11). doi:10.1083/jcb.202109133

Jhelum, P., Santos-Nogueira, E., Teo, W., Haumont, A., Lenoel, I., Stys, P. K., & David, S. (2020). Ferroptosis Mediates Cuprizone-Induced Loss of Oligodendrocytes and Demyelination. J Neurosci, 40(48), 9327–9341. doi:10.1523/JNEUROSCI.1749-20.2020

Maeda, R., Seki, N., Uwamino, Y., Wakui, M., Nakagama, Y., Kido, Y., … Sugiura, Y. (2023). Amino acid catabolite markers for early prognostication of pneumonia in patients with COVID-19. Nat Commun, 14(1), 8469. doi:10.1038/s41467-023-44266-z

Mayer, A. L., Higgins, C. B., Heitmeier, M. R., Kraft, T. E., Qian, X., Crowley, J. R., … DeBosch, B. J. (2016). SLC2A8 (GLUT8) is a mammalian trehalose transporter required for trehalose-induced autophagy. Sci Rep, 6, 38586. doi:10.1038/srep38586

Meng, Y., Heybrock, S., Neculai, D., & Saftig, P. (2020). Cholesterol Handling in Lysosomes and Beyond. Trends Cell Biol, 30(6), 452–466. doi:10.1016/j.tcb.2020.02.007

Ogretmen, B. (2018). Sphingolipid metabolism in cancer signalling and therapy. Nat Rev Cancer, 18(1), 33–50. doi:10.1038/nrc.2017.96

Perera, R. M., & Zoncu, R. (2016). The Lysosome as a Regulatory Hub. Annu Rev Cell Dev Biol, 32, 223–253. doi:10.1146/annurev-cellbio-111315-125125

Prolo, L. M., Vogel, H., & Reimer, R. J. (2009). The lysosomal sialic acid transporter sialin is required for normal CNS myelination. J Neurosci, 29(49), 15355–15365. doi:10.1523/JNEUROSCI.3005-09.2009

Rudnik, S., & Damme, M. (2021). The lysosomal membrane-export of metabolites and beyond. FEBS J, 288(14), 4168–4182. doi:10.1111/febs.15602

Ruzicka, J., Mchale, K. & Peake, D. A.. (2014). Data acquisition parameters optimization of quadrupole orbitrap for global lipidomics on LC-MS/MS time frame..

Scharenberg, S. G., Dong, W., Ghoochani, A., Nyame, K., Levin-Konigsberg, R., Krishnan, A. R., … Abu-Remaileh, M. (2023). An SPNS1-dependent lysosomal lipid transport pathway that enables cell survival under choline limitation. Sci Adv, 9(16), eadf8966. doi:10.1126/sciadv.adf8966

Thelen, A. M., & Zoncu, R. (2017). Emerging Roles for the Lysosome in Lipid Metabolism. Trends Cell Biol, 27(11), 833–850. doi:10.1016/j.tcb.2017.07.006

Traka, M., Podojil, J. R., McCarthy, D. P., Miller, S. D., & Popko, B. (2016). Oligodendrocyte death results in immune-mediated CNS demyelination. Nat Neurosci, 19(1), 65–74. doi:10.1038/nn.4193

Ueno, T., Muno, D., & Kominami, E. (1991). Membrane Markers of Endoplasmic-Reticulum Preserved in Autophagic Vacuolar Membranes Isolated from Leupeptin-Administered Rat-Liver. Journal of Biological Chemistry, 266(28), 18995–18999. Retrieved from <Go to ISI>://WOS:A1991GJ47200089

van Veen, S., Martin, S., Van den Haute, C., Benoy, V., Lyons, J., Vanhoutte, R., … Vangheluwe, P. (2020). ATP13A2 deficiency disrupts lysosomal polyamine export. Nature, 578(7795), 419–424. doi:10.1038/s41586-020-1968-7

Vasanthakumar, T., & Rubinstein, J. L. (2020). Structure and Roles of V-type ATPases. Trends Biochem Sci, 45(4), 295–307. doi:10.1016/j.tibs.2019.12.007

Wyant, G. A., Abu-Remaileh, M., Wolfson, R. L., Chen, W. W., Freinkman, E., Danai, L. V., … Sabatini, D. M. (2017). mTORC1 Activator SLC38A9 Is Required to Efflux Essential Amino Acids from Lysosomes and Use Protein as a Nutrient. Cell, 171(3), 642–654 e612. doi:10.1016/j.cell.2017.09.046

